# Spirometric indices in primary ciliary dyskinesia: systematic review and meta-analysis

**DOI:** 10.1101/486837

**Authors:** Florian S. Halbeisen, Anu Jose, Carmen de Jong, Sylvia Nyilas, Philipp Latzin, Claudia E. Kuehni, Myrofora Goutaki

**Author notes:** **Correspondence:** Myrofora Goutaki, Institute of Social and Preventive Medicine, University of Bern, Finkenhubelweg 11, 3012 Bern, Switzerland. **Take home message:** Spirometric indices of PCD patients vary between published studies, which not only suggests the possibility of methodological differences between centres, but also real differences in disease expression based on genotype-phenotype associations.

## Abstract

Primary ciliary dyskinesia (PCD) is a genetic, heterogeneous disease caused by dysfunction of cilia. Evidence is sparse and reports of lung function of PCD patients ranges from normal to severe impairment. This systematic review and meta-analysis of studies of lung function of PCD patients examines the spirometric indices of PCD patients and differences by age group and sex.

We searched PubMed, Embase, and Scopus for studies that described lung function in ≥10 patients with PCD. We performed meta-analyses and metaregression to explain heterogeneity. We included 24 studies, ranging from 13-158 patients per study. The most commonly reported spirometric indices were forced expiratory volume in 1 sec (FEV1) and forced vital capacity (FVC) presented as mean and standard deviation of percent of predicted values. We found considerable heterogeneity for both parameters (I^2^ range 94-96%). The heterogeneity remained when we stratified the analysis by age; however, FEV1 in adult patients was lower. Even after taking into account explanatory factors, the largest part of the between-studies variance remained unexplained. Heterogeneity could be explained by genetic differences between study populations, methodological factors related to the variability of study inclusion criteria, or details on the performance and evaluation of lung function measurements that we could not account for. Prospective studies therefore need to use standardised protocols and international reference values. These results underline the possibility of distinct PCD phenotypes as in other chronic respiratory diseases. Detailed characterisation of these phenotypes and related genotypes is needed in order to better understand the natural history of PCD.

## Background

Primary ciliary dyskinesia (PCD) is a genetic, heterogeneous disease caused by absence and/or dysfunction of cilia [1]. Its prevalence is estimated to be around 1 in 10000, but it is underdiagnosed [2]. It can affect ciliated cells throughout the body and lead to numerous manifestations, primarily in the respiratory system [3]. Most patients suffer from chronic and recurrent upper respiratory infections from early in life, which can lead to lung damage and bronchiectasis as the disease progresses [1,4].

Lung function is one of the most important predictors of overall survival among both healthy individuals and patients with chronic lung disease.[5] Spirometry, which is widely available, is the easiest and most common method of testing pulmonary function. Spirometric indices, primarily forced expiratory volume in 1 sec (FEV1), are used to monitor progression and describe severity in chronic respiratory diseases such as cystic fibrosis or COPD [6–8]. In PCD patients, several studies have reported results of lung function measurements, however their findings are contradictory, ranging from normal lung function to significant impairment. Most studies have focused on paediatric or young adult patients and reports from older patients are rare. Data on sex differences are also scarce, although evidence is growing that a number of pulmonary diseases affect women and men differently [9]. A detailed overview and comparison of existing studies of lung function is necessary to inform PCD research about existing and improved use of spirometry, and to determine where more evidence is needed in the study and treatment of PCD.

This systematic review and meta-analysis therefore identified all published studies presenting lung function in patients with PCD, and it summarizes spirometric indices reported in those studies. We also identify differences in these spirometric indices between male and female and between paediatric and adult patients.

## Methods

The protocol we developed for the systematic review is summarized here. We followed the PRISMA guidelines for reporting[10].

### Search strategy

We searched PubMed, Embase, and Scopus with no restriction of language or study design for studies published between January 1980 and February 2017 that describe lung function in patients with PCD (ORPHA: 244).

We tested multiple search terms to decide on the most appropriate term for each database. We selected the most concise terms that identified the largest number of studies:

Pubmed: ((“kartagener syndrome”[Title/Abstract] OR “primary ciliary dyskinesia”[Title/Abstract] OR “ciliary motility disorders”[MeSH Terms]) AND (“lung function” OR “respiratory function” OR “pulmonary function” OR “lung volume” OR “forced vital capacity” OR “forced expiratory volume” OR “FEV” OR “FEV1” OR “spirometry” OR “wash out” OR “Follow-Up Studies” [MeSH] OR “Respiratory Function Tests” [MeSH]))

Embase: ‘primary ciliary dyskinesia’:ab,ti OR ‘kartagener’:ab,ti OR ‘kartageners’:ab,ti OR ‘ciliary motility disorder’:ab,ti AND (‘lung function’ OR ‘respiratory function’ OR ‘pulmonary function’ OR ‘lung volume’ OR ‘spirometry’ OR ‘fev’ OR ‘forced expiratory volume’ OR ‘forced vital capacity’ OR ‘fev1’ OR ‘plethysmography’ OR ‘wash out’) Scopus: ((TITLE-ABS-KEY(“primary ciliary dyskinesia”)) OR (TITLE-ABS-KEY(“kartagener”)) OR (TITLE-ABS-KEY(“ciliary motility disorder”))) AND ((“wash out”) OR ((“lung function”) OR (“lung volume”) OR (“respiratory function”) OR (“pulmonary function”) OR (“forced vital capacity” OR fvc) OR (“forced expiratory volume”) OR (“spirometry”)))

After identifying all eligible studies, we checked for additional citations in their reference lists. We used the Endnote X5 (Thomson Reuters, Philadelphia, PA, USA) citation manager.

#### Definition of PCD patients

The diagnosis of PCD has changed over the period from 1980 to today [11,12]. The studies we included have a wide range of patient inclusion criteria ranging from patients with a clinical diagnosis (in most cases Kartagener syndrome) to those with positive results from tests that have varied with diagnostic consensus at the time of each study: electron microscopy (EM), light or high-speed video-microscopy (VM), nasal nitric oxide (nNO), and genetic testing.

#### Study selection

We included studies reporting lung function of patients with PCD that had a study population of 10 or more persons. We excluded animal studies and those that were not original, studies that described diagnostic test results or genetics but lacked information on lung function, and those that reported on other rare ciliary syndromes such as Joubert syndrome.

We initially screened publication titles and abstracts. From our experience with PCD literature [3], we realized that many studies of PCD patients report lung function measurements without explicitly articulating this in the title or abstract. Thus we also screened the full text of original clinical studies of PCD patients even if lung function was not mentioned in the title or abstract. After reading the full text of all potentially eligible studies, the final decision on inclusion was made by two independent reviewers, one with extensive experience in systematic reviews and knowledge of the disease. In case of disagreement, a consensus decision was reached after discussion. During the final step of inclusion, we excluded studies that did not report any spirometric indices and conference abstracts that lacked full text.

#### Overlapping study population

To avoid including the same patients multiple times in our review, we identified all studies that appeared to share the same study population. We compared the author list, country of origin, and department where the study took place. In case we noticed a considerable overlap in the study population between two or more studies, we always included in the quantitative synthesis the study that was published most recently and included information on a larger number of patients. When two studies were published 10 or more years apart, we included both since we believed there would be little chance of significant overlap. Where the extent of possible overlap was not clear, we contacted the study investigators for clarification.

#### Data extraction

We used Epidata 3.1 to extract information from all studies, including those with overlapping populations. We extracted 1) author and publication-specific information, which included author names, journal and year of publication, country and centre of corresponding author; 2) the study characteristics including years the study was carried out, study design, inclusion and exclusion criteria, study population size, country where the study took place, type of clinic, age of participants, and age stratification of reported lung function measurements; and 3) information on spirometry of PCD patients. For spirometry, we extracted all information reported on lung function measurements including values of spirometric indices and information on equipment used for spirometry, quality control measures, and reference values.

#### Meta-analysis

We performed meta-analyses of forced expiratory volume during the first second (FEV1) and forced vital capacity (FVC), the most commonly reported spirometry indices, using a DerSimonian-Laird random effects model [13,14], and assessed the heterogeneity (I^2^) between studies [15]. We assessed possible differences in lung function between children and adults by performing separate subgroup meta-analyses in adults and children. Studies that did not report FEV_1_ and FVC separately for adults and children or in which information on age was not available were excluded from this subgroup analysis.

We then investigated reasons for heterogeneity by fitting meta-regression models. We considered the following explanatory factors one at a time:

- Period of publication: before 1997, 1997–2007 and 2007–2017
- Number of patients included in the study: <20, 21–50, 51–100, and >100 patients
- Retrospective or prospective study design
- Reference values used for spirometry: references using ≥5000 subjects,<5000 subjects, no information) [16]
- Pulmonary function test quality control: based on ERS/ATS guidelines, used the best out of three valid measurements, no information available
- Whether spirometry was the main study outcome:(no, yes) [17].

Statistical analysis was performed with R software version 3.1.2 (www.r-project.org) using the metapackage (version 4.3-2), and specifically the commands metagen and metareg.

## Results

### Search

After excluding duplicates appearing in more than one of the databases (Pubmed, Embase, Scopus), our initial search produced 664 articles (Figure 1). We screened titles and abstracts of these and excluded 501 articles. Among the remaining 163 studies, 37 were either conference abstracts only or it was not possible to find the full text. After reading the full text of the remaining 126 studies, we excluded another 102 articles (Figure 1), including 34 that had overlapping study populations [18–50]. We finally included 24 studies reporting spirometric indices of PCD patients in the meta-analysis.

**Fig 1.**
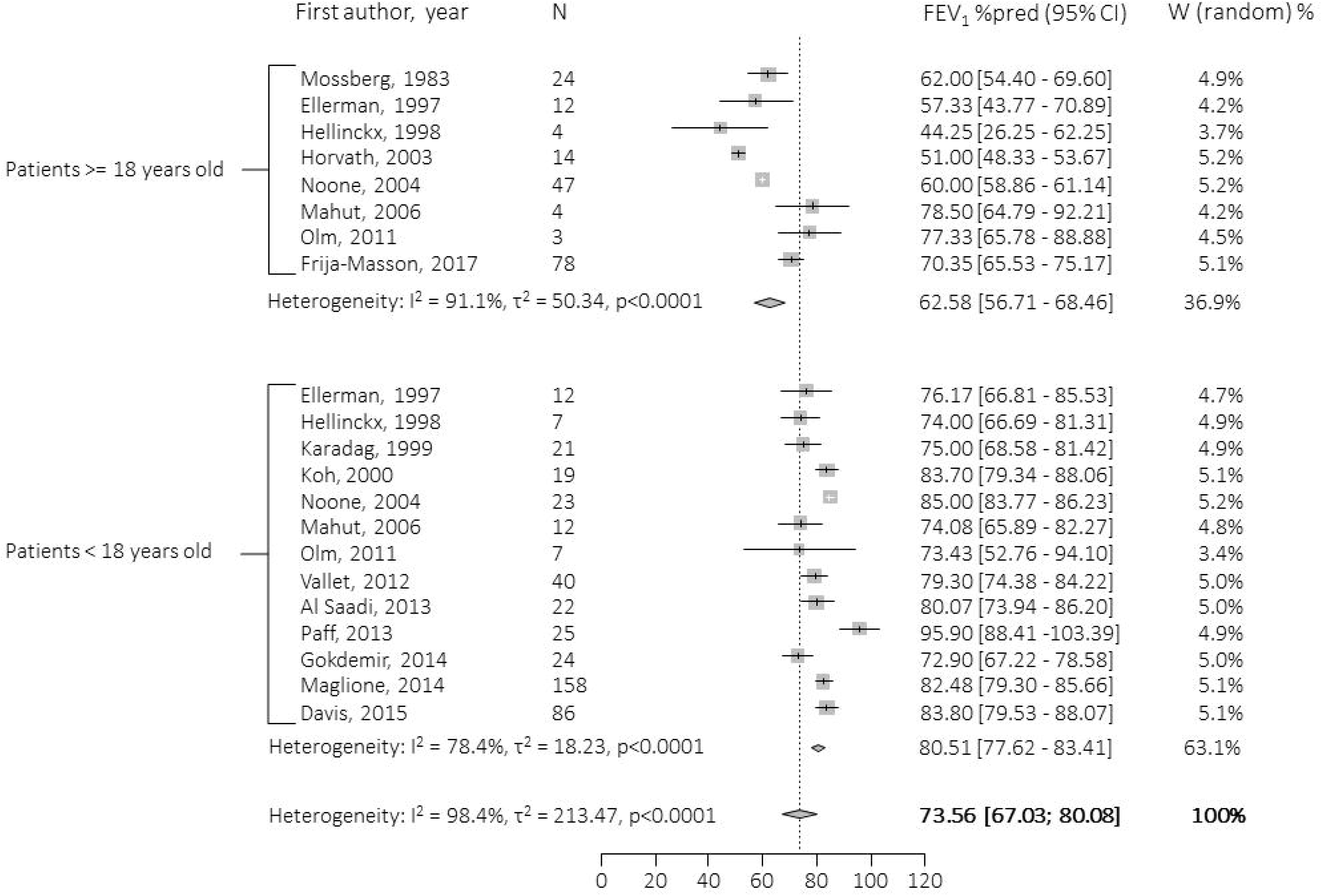
Flow chart describing the selection procedure. Data are presented as n.

### Study characteristics and information on spirometry

Table 1 lists the 24 studies and their characteristics. The articles [4,51–73] encompassed a total of 1179 patients with a mean number of 49 patients (range 13–158) per study. Most studies were relatively small, with 18 of the 24 including fewer than 50 patients (Table 2). Nearly all studies originated from paediatric or adult pulmonology departments, or from specialised PCD centres. Seventeen were single-centre studies. Fifteen were published in the last 10 years (≥2008). Fifteen came from Europe, six from Asia, and three were from North or South America. Seven studies included only children (<18 years), three only adults (≥18 years), and 14 studies had populations of mixed age consisting mainly of children with a few adults.

**TABLE 1:** Detailed characteristics of included studies describing spirometric indices of PCD patients, stratified by age group of participants

| Study | Publication Country | Publication Year | Participants n | Age | Type of study | Study design | Diagnoses | Symptoms stratified by age | Longitudinal data | FEV<sub>1</sub> | FVC |
| --- | --- | --- | --- | --- | --- | --- | --- | --- | --- | --- | --- |
| **Children** |  |  |  |  |  |  |  |  |  |  |  |
| AL SAADI et al. | S. Arabia | 2013 | 22 | 10.86±3.09<sup>#</sup> | P | Case-control | EM+HFVM | NA | - | + | - |
| DAVIS et al. | USA/Canada | 2015 | 118 | 8 (5-11)<sup>¶</sup> | P | Case series | EM or gen | NA | - | + | - |
| GOKDEMIR et al. | Turkey | 2014 | 24 | 12.9±2.7<sup>#</sup> | P | Clinical trial | EM or Kartagener | NA | - | + | + |
| KARADAG et al. | UK | 1999 | 21 | 10.8±3.2<sup>#</sup> | R | Case-control | EM, HFVM | NA | - | + | + |
| KOH et al. | Korea | 2000 | 19 | 12 (7-16)<sup>+</sup> | P | Clinical trial | EM | NA | - | + | - |
| MAGLIONE et al. | Denmark, Italy, UK | 2014 | 158 | 8.7 (4.2–17.4) | R | Cohort | according to consensus | NA | + | + | + |
| PAFF et al. | The Netherlands | 2013 | 25 | 10.7 (7.1–14.5)<sup>¶</sup> | P | Case-control | according to consensus | NA | - | + | - |
| **Adults** |  |  |  |  |  |  |  |  |  |  |  |
| FRIJA-MASSON et al. | France | 2017 | 78 | 34.8 (28.6–47.1)<sup>¶</sup> | R | Case series | EM, gen or Kartagener | NA | + | + | + |
| HORVATH et al. | UK | 2003 | 14 | 35±4.6<sup>#</sup> | P | Case-control | EM, HFVM | NA | - | + | + |
| MOSSBERG et al. | Sweden | 1983 | 24 | 30 (19-47)<sup>+</sup> | P | Case series | clinical suspicion | NA | - | + | - |
| **Both children and adults** |  |  |  |  |  |  |  |  |  |  |  |
| AMIRAV et al. | Israel | 2016 | 13 | 10 (5-43)<sup>¶</sup> | R | Case series | all tests, including gen | - | - | + | - |
| BOON et al. | Belgium | 2014 | 122 | 17.7 (9.5–28.1)<sup>¶</sup> | R | Case series | EM, HFVM, gen | - | - | + | + |
| COHEN-CYMBERKNOH et al. | Israel | 2014 | 14 | 15.9±8.6<sup>#</sup> | R | Case series | EM, HFVM, nNO, gen | - | - | + | - |
| ELLERMAN et al. | Denmark | 1997 | 24 | 21±13.6<sup>#</sup> | P | Case series | HFVM | + | + | + | + |
| HELLINCKX et al. | Belgium | 1998 | 11 | 15.2±7<sup>#</sup> | P | Case series | EM, HFVM | + | + | + | - |
| MAHUT et al. | France | 2006 | 16 | 13.5 (12-17)<sup>¶</sup> | R | Case series | Clinical, EM | + | - | + | - |
| NOONE et al. | USA/Canada | 2004 | 78 | In adults: 36 (19-73)<sup>+</sup><br>In children: 8 (1-17)<sup>+</sup> | P | Case series | EM, HFVM, gen | + | - | + | - |
| NYILAS et al. | Germany/Switzerland | 2017 | 49 | 14.7±6.6<sup>#</sup> | P | Case-control | all tests | - | - | + | + |
| OLM et al. | Brazil | 2011 | 1045 | 13.4±4.7<sup>#</sup> | P | Case-control | EM+HFVM | + | - | + | + |
| PIFFERI et al. | Italy | 2014 | 45 | 14 (22.25)<sup>¶</sup> | P | Case-control | EM, HFVM, nNO | - | - | + | + |
| SHAH et al. | UK | 2016 | 118 | 23.5 (10-36)<sup>¶</sup> | R | Case series | EM, HFVM, nNO, 2% clinical | - | + | + | - |
| TOLUSAKOW et al. | Germany | 1981 | 21 | NR (3-43)<sup>+</sup> | R | Case series | clinical suspicion | - | - | + | + |
| VALLET et al. | France | 2013 | 41 | 6 (3-11)<sup>¶</sup> | R | Case series | EM, HFVM | - | + | + | + |
| YIALLOUROS et al. | Cyprus | 2015 | 20 | NR (0.1-58.4)<sup>+</sup> | R | Case series | EM or HFVM+nNO | + | + | + | + |

Age is reported in years as mean ±SD<sup>#</sup>, mean (range)<sup>+</sup>, median (IQR)<sup>¶</sup>, or median (range)<sup>+</sup> depending on the available information; NR, not reported R, retrospective; P, prospective; EM, electron microscopy; HFVM, high frequency video-microscopy; nNO, nasal NO; gen, genetic analysis; NA, not applicable

PCD diagnosis was established in different ways. Nineteen studies assessed ciliary ultrastructure using electron microscopy and/or a combination of one or more of nasal nitric oxide, video microscopy, and genetics according to available consensus at the time of each publication (Table 2). In three of the 24 studies, the majority of patients were similarly diagnosed on the basis of ciliary ultrastructure, though a small proportion of the patients were diagnosed based only on strong clinical suspicion—mainly Kartagener syndrome. In two, older studies, all patients were diagnosed based on strong clinical suspicion, again, primarily Kartagener syndrome.

**Table 2:** Characteristics of included studies describing spirometry indices of PCD patients

| **Characteristic** | **N (%)** |
| --- | --- |
| **Total studies** | 24 (100) |
| **Study size** |  |
| ≤20 patients | 8 (33) |
| 21-50 patients | 10 (42) |
| 50-100 patients | 2 (8) |
| >100 patients | 4 (17) |
| **Number of centres** |  |
| Single centre | 17 (71) |
| Multicentre | 7 (29) |
| **Publication period** |  |
| ≤1996 | 2 (8) |
| 1997-2007 | 7 (29) |
| ≥2008 | 15 (63) |
| **Study region** |  |
| Europe | 15 (62.5) |
| Asia | 6 (25) |
| North and South America | 3 (12.5) |
| **Age of participants** |  |
| Children <18 years | 7 (29) |
| Adults ≥18 years | 3 (13) |
| Children and adults | 14 (58) |
| **Study design** |  |
| Retrospective | 11 (46) |
| Prospective | 13 (54) |
| **Diagnostics performed** |  |
| Diagnosis based only on strong clinical suspicion | 2 (8) |
| Diagnosis based on EM or a combination of tests in most patients; few only on strong clinical suspicion¶ | 3 (13) |
| Diagnosis based on EM or a combination of tests in all patients¶ | 19 (79) |
| **Spirometry indices reported** |  |
| FEV1 | 24 (100) |
| FVC | 12 (50) |
| **Information for different age groups** | 6<sup>#</sup> (43) |
| **Longitudinal information** | 8 (33) |

¶ Combination of the following: electron microscopy, nasal NO, high frequency videomicroscopy or light microscopy, genetics

All studies reported FEV1 measurements and half of them also reported FVC measurements. How investigators reported lung function measurements varied. The most commonly reported spirometric indices were FEV1 and FVC, presented as mean and standard deviation (SD) of percent predicted values. In studies in which FEV1 and FVC were reported differently or important information was not reported, we contacted the investigators for additional data. Among the 14 studies that included both paediatric and adult patients, only six of the 14 reported lung function results stratified by age groups. Only four of the 24 studies reported FEV1 and FVC as z-scores. Eight also reported other lung function measurements such as plethysmography and multiple breathe washout techniques, or exhaled breath analysis techniques. Investigators used a variety of different spirometry reference values depending on the year of publication; 10 out of the 24 studies did not report which references were used for the calculation of the percent predicted values. Twelve of the studies reported that spirometry was performed based on the ERS/ATS guidelines, while in five more the best of three acceptable spirometry measurements was chosen. The remaining seven studies did not report on quality measures of spirometry. Only seven of the 24 studies included longitudinal measurements of lung function.

Table S1 presents the characteristics of all included studies and studies with overlapping population.

### FEV1

Reported FEV1 values varied significantly between the studies and mean FEV1 %predicted ranged from 51% to 96%. Weighted mean FEV1 in the 24 studies was 75% (95% C.I. 69–80%) with a heterogeneity of I^2^= 96% (Figure 2). Results remained the same when we excluded from the meta-analysis the two studies having only clinically diagnosed patients. In the subgroup meta-analyses, including the 16 studies with information on age groups, eight studies reported FEV1 measurements in patients ≥18 years old and 13 reported FEV1 measurements in children younger than 18 years. In adults, mean FEV1 percent predicted values ranged from 44% to 79% with a weighted mean value of 63% (95% C.I. 57–69%, Figure 3) and I^2^=91. In children, mean FEV1 percent predicted values ranged from 73% to 85% with a weighted mean value of 81% (95%C.I. 78–83%; Figure 3) and I^2^=78.

**Fig 2.**
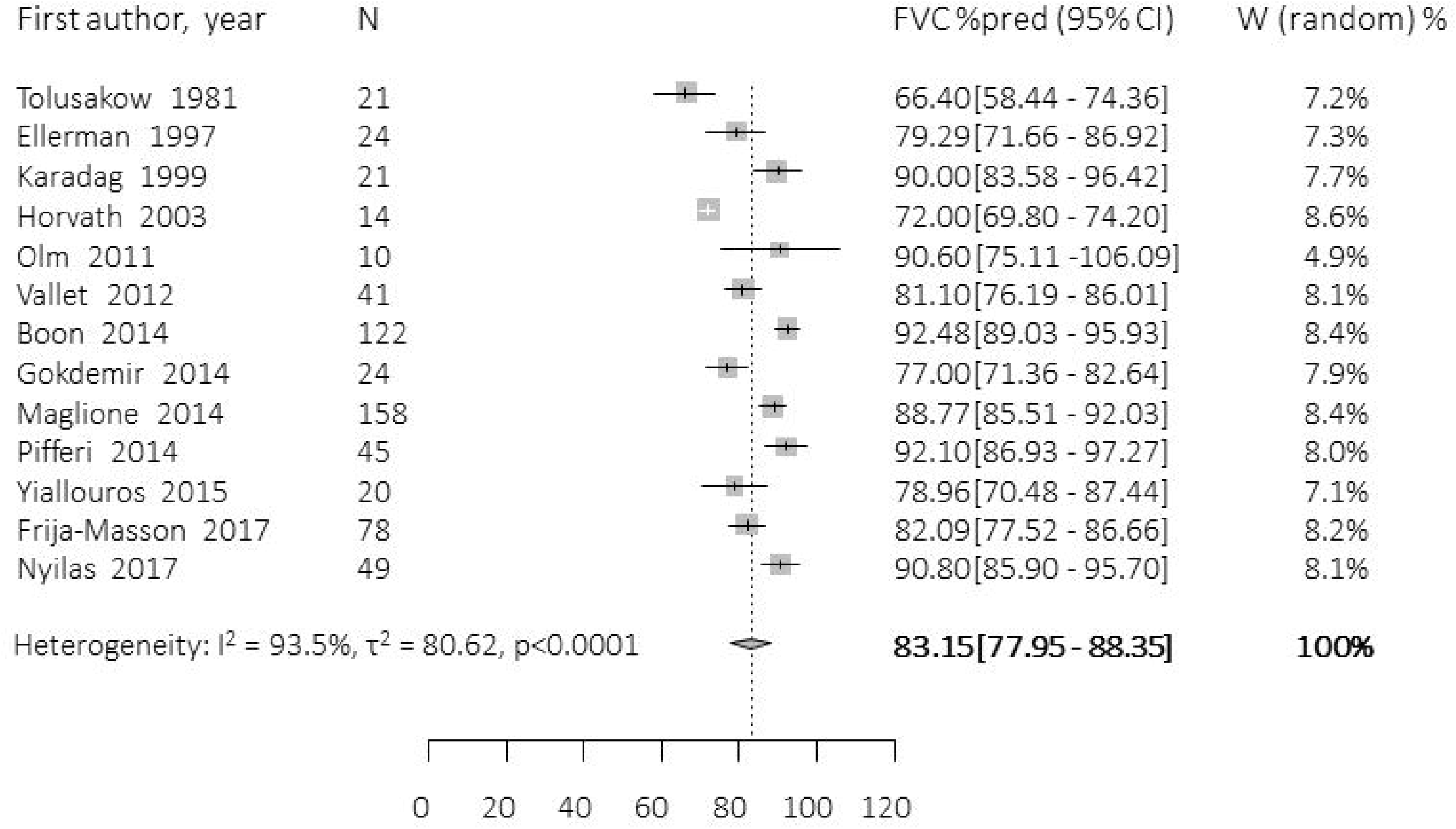
FEV1 in primary ciliary dyskinesia patients. forest plot showing the heterogeneity and weighted mean value of FEV1 percent predicted in the included publications.

**Fig 3.**
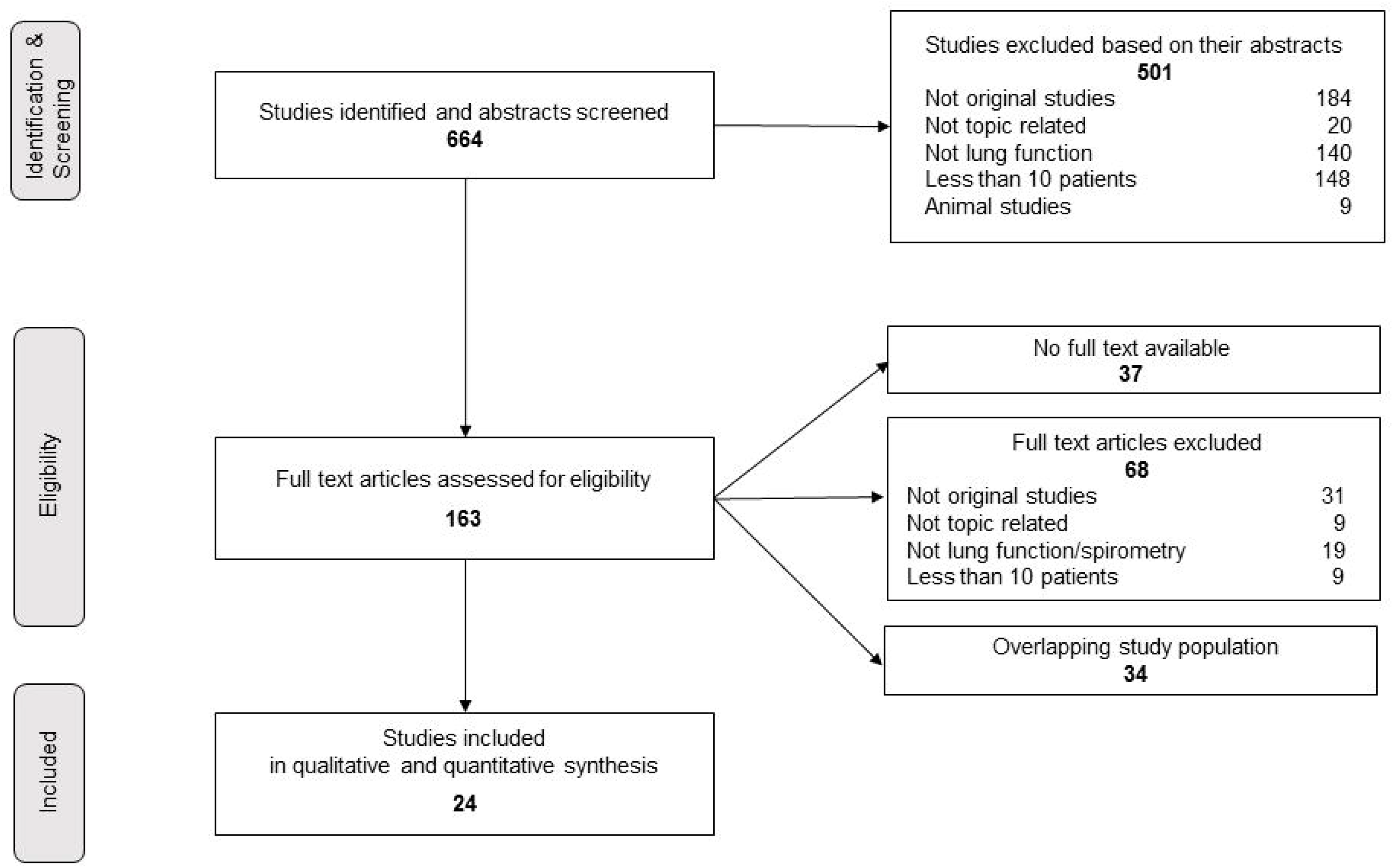
FEV1 in primary ciliary dyskinesia patients by age group. forest plot showing the heterogeneity and weighted mean value of FEV1 percent predicted in the included publications.

### FVC

In the 13 studies including FVC results, reported values ranged from 66% to 92%. Weighted mean FVC was 83% (95% C.I. 69–80%) with a heterogeneity of I^2^= 94% (Figure 4). In the subgroup meta-analyses, we included four studies reporting FVC measurements for patients ≥18 years old and six studies of children younger than 18 years. The mean FVC percent predicted values in adults ranged from 70% to 94% with a weighted mean value of 79% (95% C.I. 70–88%, figure S1) and I^2^=89%. In children, the mean FVC percent predicted ranged from 77% to 90% with a weighted mean value of 85% (95%C.I. 80–90%; Figure 2) and I^2^=72.

**Fig 4.**
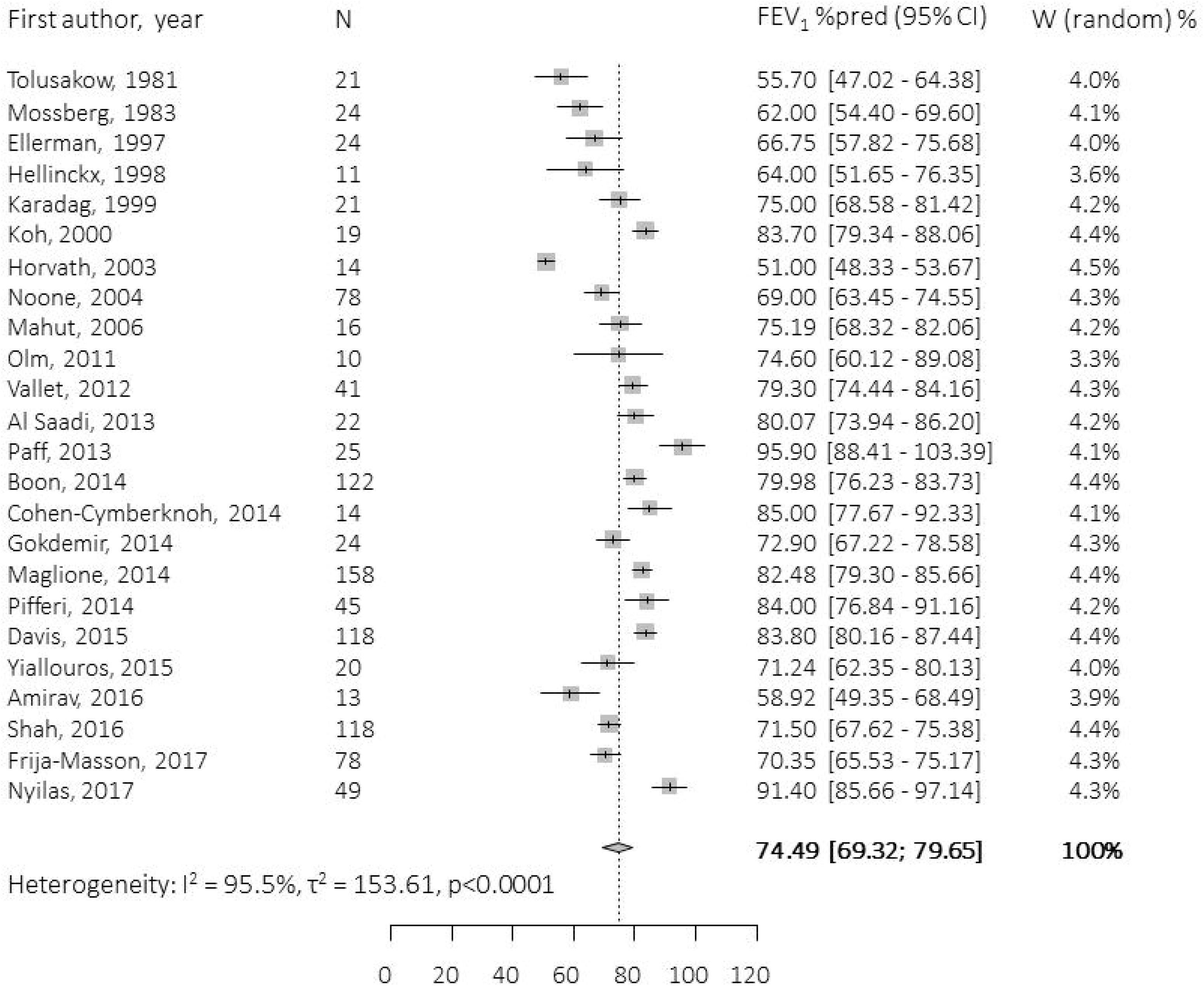
FVC in primary ciliary dyskinesia patients. forest plot showing the heterogeneity and weighted mean value of FVC percent predicted in the included publications.

### Meta-regression

Meta-regressions showed that the explanatory factors explained only a small part of the between-studies variance for all symptoms. Studies published after 2007 reported higher mean values of FEV1 and FVC (both p=0.01). Another factor that explained part of the heterogeneity was the reference values for spirometry. Studies that did not report the reference values they used reported lower mean values for both FEV_1_ and FVC (p=0.003 and 0.002, respectively). Mean FEV_1_ was lower in studies that did not include any information on pulmonary function test quality control (p=0.02). Detailed results of the meta-regression models are in table S2.

### Differences between males and females

Only two of the 24 studies compared lung function of male and female patients. In a large cohort of French adult patients, the spirometric indices of women were lower than those of men at the end of the study, and the annual decline of FEV1 was also greater in women [57]. In contrast, no difference by sex was reported in a large British adult cohort [70].

### Longitudinal data

Finally, though a preponderance of PCD studies made cross-sectional assessments, seven of the 24 studies in our meta-analysis followed lung function of PCD patients longitudinally, as did a further five of the 34 studies in which we observed overlapping populations. The differing designs and follow-up times of these 12 longitudinal studies disallow pooling their results in a meta-analysis. However, we offer a qualitative overview of the findings reported in them.

Four studies reported that most patients had stable spirometric indices (FEV1, FVC) during respective short-term or longer-follow-ups [56,59,63,72]. Four other, more recent studies including adult patients reported important declines of FEV1 and/or FVC [27,40,57,70]. A study of the Danish national cohort of PCD patients [25] reported at least a 10% decrease in FEV1 percent predicted in one-third of the patients, though great variation in lung function was observed after follow-up. In the British adult cohort, ultrastructural defect was associated with lung function decline: patients with microtubular defect reported a steeper decline [70], a finding that was not consistent with the results of a study of 60 French children with PCD [72]. Larger national or multicentre studies reported no association between lung function decline and age of diagnosis [25,63]. Preliminary results from 71 patients included in the international PCD registry showed an annual mean decline of 0.59% in percent predicted FEV1 and of 0.39% in percent predicted FVC [40]. Recent, smaller studies have evaluated the effect of managing lung function of PCD patients with, for example, lobectomy [73] and endoscopic sinus surgery [18] without observing significant differences in lung function decline.

## Discussion

This is the first systematic review of lung function measurements of PCD patients. FEV1 was lower in adults than children, but studies report a wide range of FEV1 and FVC values. This heterogeneity is not accounted for by available explanatory factors. Of the 664 studies originally identified, only 9% reported spirometry of PCD patients; 34 of the studies had overlapping study populations. Most studies were small, with a mean of 49 patients per study, and fewer than half of the studies that included both children and adults reported lung function stratified by age. Only two studies examined differences by sex. Reports of spirometric indices were associated with the period of publication, provision of lung function reference values, and information on spirometry quality control. Not surprisingly, newer studies reported higher lung function values.

The main strength of this study inheres in its methodology. We performed the search without language restrictions and screened the full text of every identified clinical study with an original study population—even if lung function was not explicitly mentioned in the abstract. This ensured that we included studies that a custom approach might miss. We contacted primary investigators of the included studies to acquire spirometric indices in a consistent form in order to pool them in meta-analyses, and we separated our results for paediatric and adult patients. We also accounted for heterogeneity in the results by performing a metaregression with all available explanatory factors. However, we were not able to include other factors that were possibly known but not published in the studies. These include genetic variations in studied populations, which are known to result in varying disease severity [52,55], other indicators of disease severity such as imaging data to assess structural changes in the lungs, and less well documented clinical factors such as compliance, adherence to therapy, age at diagnosis, and varying standards in clinical surveillance and treatment (e.g. history of lobectomy, clinical status at the time of measurement).

We restricted our search to studies published since 1980 for several reasons. Many things have changed in understanding and characterising PCD since it was first described by Kartagener in early 1930s. Older studies are often not available online, or do not have an abstract in the online databases. In addition, clinical studies of PCD before 1980 were mostly case reports or small studies with fewer than 10 patients that were not eligible for our review.

Our review undoubtedly reflects the weaknesses of its included studies, most of which suffered from significant selection bias. Its important limitations involve the differences in study designs and aims of the included studies that might have directly influenced the lung function assessment of study populations. We did not perform a risk of bias assessment for the studies included in this review. We expected all studies to suffer from moderate to serious risk, which is common in this type of studies, especially in rare diseases e.g. small studies mainly based on case series of patients. Our most important limitation is that we could not compare the spirometric indices of the included studies using the same reference values and, ideally, use z-scores for FEV1 and FVC. Unfortunately, the majority of included studies reported percent predicted values derived from a variety of references. To address this issue, we tested the particular lung function reference as a possible explanatory factor in the heterogeneity in our metaregression.

We do not expect significant diagnostic misclassification in the included studies, mainly because most studies adhered to consensus guidelines and were published more recently. However, misclassification bias is still possible, especially for the older studies. When we repeated our meta-analyses excluding the two studies with only clinical diagnoses the results did not change. Because most studies suffered from the same design flaws, we did not apply any quality assessment criteria to decide which studies to include in our meta-analysis.

Due to the considerable heterogeneity, the calculated mean weighted values of percent predicted FEV1 and FVC should be interpreted with caution. The meta-analysis was performed to quantify the variability in reported spirometric indices and not to give valid estimates. The possible explanatory factors we tested failed to explain this heterogeneity. Still, some factors contributed to explaining differences in FEV1 and FVC in the included studies. Publication year may be indicative of differences in awareness and disease diagnosis as well as changes in standardisation of treatment and routine surveillance of patients. Recent studies are more likely to have used modern diagnostic approaches and include more patients who are correctly diagnosed and patients who have less severe disease, and more genetic variations compared to older studies. They also usually include patients treated with more modern treatment protocols and use newer and more correct spirometry reference values. Studies that did not report information on quality control of spirometry data or lung function reference values might have had less strict protocols for lung function measurements compared to studies that reported this information in detail. Although it was not possible to pool the studies including repeated lung function measurements and study disease progression over time, the reported percent predicted FEV1 was lower in adult patients compared to children. On one hand, this could support the progression of lung disease into adulthood; on the other hand, differences between adults and children might also reflect differences in disease management or diagnosis over the years.

Methodological variability in the studies we included could not explain the heterogeneity in lung function. We believe that this heterogeneity is caused by factors that could not be tested in a meta-analysis given the information that we had available (such as genetic factors, differences in disease management, factors related to the variability of study inclusion criteria, and details on the performance and evaluation of lung function measurements). This heterogeneity in lung function measurements goes hand in hand with the heterogeneity we reported in our previous review on the clinical manifestations of PCD, which highlights the limitations of published studies to describe the full clinical spectrum of PCD [3]. However, the results of both reviews underline the possibility of distinctly different disease phenotypes, which also are observed in other chronic respiratory diseases. This is further supported by reports that link specific ultrastructural defects and genetic mutations with more severe lung disease. In a study of North American children with PCD, those with an absence of inner dynein arms in conjunction with central apparatus defects and microtubular disorganisation associated with CCDC39 and CCDC40 mutations had lower spirometric indices compared to other PCD patients [55]. In a smaller case series of Israeli patients, in combination with previous case reports, patients with CCNO mutations showed rapid deterioration of lung function [52,74,75].

All these findings demonstrate the need for collaborative, prospective studies that use standardised protocols for spirometry, and international up-to-date reference values. To meet this need, together with a network of international collaborators we are developing a standardised, disease-specific instrument for patient follow-up and prospective data collection [76,77]. At the same time, researchers in centers around the world are conducting a prospective multicentre observational study of variability of lung function in PCD patients (PROVALF-PCD) that will provide information on how FEV1 itself varies in individual stable PCD patients [78].

## Conclusion

Spirometric indices of PCD patients vary significantly between published studies, which not only suggests the possibility of methodological differences between centres, but also real differences in disease expression. Detailed characterisation of different PCD phenotypes and, ideally, phenotype-genotype associations are needed to explain this variability, and better understand the natural history of PCD.

## Supporting information

## Acknowledgements

We want to thank Christopher Ritter (Institute of Social and Preventive Medicine, University of Bern, Switzerland) for his editorial suggestions.

## Authors’ contributions

MG and CK developed the concept and designed the study. MG and AJ performed the systematic search, selected the included studies and extracted the data. FH and MG analysed the data and drafted the initial manuscript. All authors contributed to iterations and approved the final version. MG, CK and FH take final responsibility for the contents.

---

**Supplementary Figure S1** FVC in primary ciliary dyskinesia patients by age group, Fig S1 legend: Forest plot showing the heterogeneity and weighted mean value of FVC percent predicted in the included publications.

