## Supplementary material for "Spirometric indices in primary ciliary dyskinesia: systematic review and meta-analysis"

TABLE S2: Metaregression results on spirometric indices (FEV1 and FVC) of PCD patients

|  | |  | **Regression coefficient** | | | **95% C.I.** | | | **p-value^¶^** | | **Residual I^2^**  **(p-value)^#^** | | | | |  | | **Regression coefficient** | | | **95% C.I.** | | | **p-value^¶^** | | **Residual I^2^**  **(p-value)^#^** |
| --- | --- | --- | --- | --- | --- | --- | --- | --- | --- | --- | --- | --- | --- | --- | --- | --- | --- | --- | --- | --- | --- | --- | --- | --- | --- | --- |
|  | |  | **FEV_1_** | | | | | | | | | |  | | |  | | **FVC** | | | | | | | | |
| **Year of publication** | **>2007*** | | | 79.0 | 73.6 | | 84.3 | <.0001 | | 93.03% (< .0001) | | | |  | 86.0 | | | | | 81.4 | | 90.6 | <.0001 | | 85.99% (< .0001) | |
|  | **1997-2007** | | | -9.3 | -19.3 | | 0.7 | **0.069** | |  | |  | |  | -6.1 | | | | | -15.2 | | 2.9 | 0.183 | |  | |
|  | **≤1997** | | | -17.5 | -30.9 | | -4.1 | **0.011** | |  | |  | |  | -19.6 | | | | | -34.9 | | -4.3 | **0.012** | |  | |
| **Study size** | **≤20 patients*** | | | 70.4 | 62.0 | | 78.8 | <.0001 | | 94.12% (< .0001) | | | |  | 78.0 | | | | | 69.1 | | 86.8 | <.0001 | | 84.38% (< .0001) | |
|  | **21-50 patients** | | | 6.1 | -5.1 | | 17.2 | 0.287 | |  | |  | |  | 4.8 | | | | | -5.5 | | 15.2 | 0.361 | |  | |
|  | **51-100 patients** | | | -0.8 | -18.9 | | 17.3 | 0.934 | |  | |  | |  | 4.1 | | | | | 12.1 | | 20.3 | 0.618 | |  | |
|  | **>100 patients** | | | 9.0 | -5.0 | | 23.0 | 0.207 | |  | |  | |  | 12.7 | | | | | -0.2 | | 25.5 | 0.054 | |  | |
| **Study design** | **prospective*** | | | 75.4 | 68.3 | | 82.6 | <.0001 | | 95.50% (< .0001) | | | |  | 83.1 | | | | | 75.7 | | 90.4 | <.0001 | | 91.92% (< .0001) | |
|  | **retrospective** | | | -2.0 | -12.6 | | 8.5 | 0.703 | |  | |  | |  | 0.2 | | | | | -9.7 | | 10.1 | 0.972 | |  | |
| **Spirometry references** | **≥5000 participants*** | | | 82.2 | 74.6 | | 89.9 | <.0001 | | 91.66% (< .0001) | | | |  | 90.7 | | | | | 84.0 | | 97.3 | <.0001 | | 82.70% (< .0001) | |
|  | **<5000 participants** | | | -4.5 | -14.9 | | 5.8 | 0.393 | |  | |  | |  | -5.5 | | | | | -14.9 | | 4.0 | 0.256 | |  | |
|  | **no information** | | | -15.0 | -24.7 | | -5.2 | **0.003** | |  | |  | |  | -12.9 | | | | | -21.3 | | -4.6 | **0.002** | |  | |
| **Lung function test quality** | **ERS/ATS guidelines*** | | | 79.9 | 73.5 | | 86.4 | <.0001 | | 93.55% (< .0001) | | | |  | 85.0 | | | | | 79.9 | | 90.2 | <.0001 | | 85.58% (< .0001) | |
|  | **best of 3 measurements** | | | -8.7 | -20.4 | | 3.0 | 0.144 | |  | |  | |  | -9.9 | | | | | -21.1 | | 1.4 | 0.085 | |  | |
|  | **no information** | | | -12.1 | -22.6 | | -1.6 | **0.024** | |  | |  | |  | -1.0 | | | | | -10.8 | | 8.7 | 0.834 | |  | |
| **Study design** | **case-control*** | | | 78.7 | 68.7 | | 88.8 | <.0001 | | 95.21% (< .0001) | | | |  | 86.7 | | | | | 77.0 | | 96.3 | <.0001 | | 93.82% (< .0001) | |
|  | **case series** | | | -7.5 | -19.8 | | 4.8 | 0.229 | |  | |  | |  | -6.3 | | | | | -19.3 | | 6.6 | 0.339 | |  | |
|  | **clinical trial** | | | 3.8 | -23.8 | | 31.3 | 0.790 | |  | |  | |  | 2.1 | | | | | -20.6 | | 24.8 | 0.856 | |  | |
|  | **cohort** | | | -0.4 | -21.3 | | 20.6 | 0.972 | |  | |  | |  | -9.7 | | | | | -32.8 | | 13.5 | 0.413 | |  | |
| **Spirometry as primary outcome** | **No*** | | | 74.2 | 68.0 | | 80.3 | <.0001 | | 95.38% (< .0001) | | | |  | 82.6 | | | | | 75.7 | | 89.5 | <.0001 | | 93.37% (< .0001) | |
|  | **Yes** | | | 1.1 | -10.3 | | 12.5 | 0.854 | |  | |  | | | | |  | | 1.4 | -9.5 | | 12.3 | 0.802 | |  | |

95% C.I. = 95% confidence interval, *reference category, ^¶^ p-value of QM test, #Residual I^2^ heterogeneity (p-value of QE test). Table shows estimated differences (95% C.I.) in percent predicted of each category of the tested moderator characteristics from the reference category. Negative (positive) values represent decrease (increase) in percent predicted values. For the reference categories, the absolute value in percent predicted is shown (intercept). The p-values are from the Q_M_ moderators’ test, which tests whether at least one of the regression coefficients (not including the intercept) is different from zero, and from the Q_E_ test for residual heterogeneity, which tests whether the heterogeneity is explained by the moderator characteristic.
