## Supplementary material for "Spirometric indices in primary ciliary dyskinesia: systematic review and meta-analysis"

**TABLE S1:** Characteristics of included studies describing spirometric indices of PCD patients by country, including studies with overlapping population

| **Study** | **Publication**  **Year** | **Participants n** | **Age** | **Type of study** | | **Study design** | **Diagnostics** | **Symptoms stratified by age** | **Longitudinal data** | **FEV1** | **FVC** |
| --- | --- | --- | --- | --- | --- | --- | --- | --- | --- | --- | --- |
| **Single-centre/regional studies** | |  |  | |  |  |  |  |  |  |  |
| **Belgium** |  |  |  | |  |  |  |  |  |  |  |
| BOON et al. | 2015 | 39 | 16.1 (11.1-19.6) ^¶^ | | P | Case-control | EM, HFVM | - | - | + | - |
| **BOON et al.** | 2014 | 122 | 17.7 (9.5–28.1)^¶^ | | R | Case series | EM, HFVM, gen | - | - | + | + |
| BOON et al. | 2014 | 11 | 16 (7-62) ^+^ | | R | Case series | EM, HFVM, gen | + | - | + | - |
| **HELLINCKX et al.** | 1998 | 11 | 15.2±7^#^ | | P | Case series | EM, HFVM | + | + | + | - |
| **Brazil** |  |  |  | |  |  |  |  |  |  |  |
| **OLM et al.** | 2011 | 104 | 13.4±4.7^#^ | | P | Case-control | EM+HFVM | + | - | + | + |
| **Canada** |  |  |  | |  |  |  |  |  |  |  |
| RATJEN et al. | 2016 | 35 | 13.7 (5.8-17.9) ^+^ | | P | Case-control | EM, gen | - | - | + | + |
| RASHAKRISHNAN ET AL | 2012 | 10 | 12.1 (7.8-15.4) ^+^ | | P | Case-control | EM, nNO, gen | - | - | + | + |
| **Cyprus** |  |  |  | |  |  |  |  |  |  |  |
| **YIALLOUROS et al.** | 2015 | 20 | NR (0.1-58.4) ^*^ | | R | Case series | EM or HFVM+nNO | + | + | + | + |
| **Denmark** |  |  |  | |  |  |  |  |  |  |  |
| ALANIN et al. | 2016 | 24 | 24 (10-65) ^+^ | | P | Case series | EM, HFVM, gen | - | + | + | + |
| GREEN et al. | 2016 | 28 | 12.4 (10.7-14.6) ^¶^ | | P | Case-control | EM, HFVM | - | - | + | + |
| LOOMBA et al. | 2016 | 17 | 13.8 ± 3.5^#^ | | R | Case-control | EM, HFVM, nNO | - | - | + | + |
| MARTHIN et al. | 2010 | 74 | 9 (4.4–43.7) ^+^ | | R | Case series | EM, HFVM, nNO | - | + | + | + |
| MADSEN et al. | 2013 | 44 | 14.8 (6.5-29.7) ^+^ | | P | Case-control | EM, HFVM, nNO | - | - | + | + |
| GREEN et al. | 2012 | 27 | 11.3 (6.3-18.5) ^+^ | | P | Case series | EM, HFVM | - | - | + | + |
| **ELLERMAN et al.** | 1997 | 24 | 21± 13.6^#^ | | P | Case series | HFVM | + | + | + | + |
| GROTH et al. | 1989 | 14 | 23.5 (12-44) ^+^ | | P | Case-control | Clinical, EM | - | - | + | + |
| **France** |  |  |  | |  |  |  |  |  |  |  |
| **FRIJA-MASSON et al.** | 2017 | 78 | 34.8 (28.6–47.1)^¶^ | | R | Case series | EM, gen or Kartagener | NA | + | + | + |
| **VALLET et al.** | 2012 | 41 | 6 (3-11)^¶^ | | R | Case series | EM, HFVM | - | + | + | + |
| MAGNIN et al. | 2012 | 20 | 7.5 (7.1-8.7) ^¶^ | | R | Case series | EM | - | + | + | + |
| **MAHUT et al**. | 2006 | 16 | 13.5 (12-17)^¶^ | | R | Case series | Clinical, EM | + | - | + | - |
| **Germany** |  |  |  | |  |  |  |  |  |  |  |
| **TOLUSAKOW et al.** | 1981 | 21 | NR (3-43) ^*^ | | R | Case series | clinical suspicion | - | - | + | + |
| **Israel** |  |  |  | |  |  |  |  |  |  |  |
| **AMIRAV et al.** | 2016 | 13 | 10 (5-43)^¶^ | | R | Case series | all tests, including gen | - | - | + | - |
| **COHEN-CYMBERKNOH et al.** | 2014 | 14 | 15.9±8.6^#^ | | R | Case series | EM, HFVM, nNO, gen | - | - | + | - |
| Age is reported in years as mean ±SD^#^, mean (range)^*^, median (IQR)^¶^, or median (range)^+^ depending on the available information; NR, not reported  R, retrospective; P, prospective; EM, electron microscopy; HFVM, high frequency video-microscopy; nNO, nasal nitric oxide; gen, genetic analysis; NA, not applicable  Studies appearing in bold are those included in the meta-analysis | | | | | | | | | | | |
| **Italy** |  |  |  | |  |  |  |  |  |  |  |
| **PIFFERI et al.** | 2014 | 45 | 14 (22.25)^¶^ | | P | Case-control | EM, HFVM, nNO | - | - | + | + |
| MIRRA et al. | 2014 | 22 | 10.5 (2-34) ^+^ | | P | Case series | EM, HFVM | - (not by age) | - | + | + |
| VALERIO et al. | 2012 | 24 | 7.6 (0.1-19) ^+^ | | P | Case series | EM, HFVM | - | - | + | + |
| PIFFERI et al. | 2012 | 50 | In adults: 30.5 (18-47) ^+^  children: 11 (6-17) ^+^ | | P | Case series | EM, HFVM | - | - | + | + |
| MAGLIONE et al. | 2012 | 20 | 11.6 (6.5-27.5) ^+^ | | R | Case series | EM, HFVM | - | - | + | + |
| MONTELLA et al. | 2009 | 13 | 15.2 (10.4-29.3) ^+^ | | P | Case series | EM, HFVM | + | - | + | + |
| SANTAMARIA et al. | 2008 | 14 | 15 (7-27) ^+^ | | P | Case-control | Clinic, EM | - | - | + | + |
| **Korea** |  |  |  | |  |  |  |  |  |  |  |
| **KOH et al**. | 2000 | 19 | 12 (7-16) ^+^ | | P | Clinical trial | EM | NA | - | + | - |
| **Saudi Arabia** |  |  |  | |  |  |  |  |  |  |  |
| **AL SAADI et al.** | 2013 | 22 | 10.86±3.09^#^ | | P | Case-control | EM+HFVM | NA | - | + | - |
| **Sweden** |  |  |  | |  |  |  |  |  |  |  |
| **MOSSBERG et al**. | 1983 | 24 | 30 (19-47)^*^ | | P | Case series | clinical suspicion | NA | - | + | - |
| **Turkey** |  |  |  | |  |  |  |  |  |  |  |
| **GOKDEMIR et al.** | 2014 | 24 | 12.9±2.7^#^ | | P | Clinical trial | EM or Kartagener | NA | - | + | + |
| **The Netherlands** |  |  |  | |  |  |  |  |  |  |  |
| **PAFF et al.** | 2013 | 25 | 10.7 (7.1–14.5)^¶^ | | P | Case-control | according to consensus | NA | - | + | - |
| **United Kingdom** |  |  |  | |  |  |  |  |  |  |  |
| **SHAH et al.** | 2016 | 118 | 23.5 (10-36)^¶^ | | R | Case series | EM, HFVM, nNO,  2% clinical | - | + | + | - |
| SUNTHER et al. | 2016 | 150 | 11.4 (6-16.2) ^+^ | | R | Case series | EM, HFVM, nNO | - | + | + | + |
| IRVING et al. | 2013 | 33 | 24.66^*^ | | P | Case-control | EM, HFVM, nNO | - | - | + | + |
| SHOEMARK et al. | 2008 | 20 | 40 (32-45) ^+^ | | P | Case-control | EM, HFVM | - | - | + | - |
| PARASKAKIS et al. | 2007 | 24 | 12 (8-16) ^+^ | | P | Case-control | EM, HFVM, nNO | - | - | + | - |
| ZIHLIF et al. | 2006 | 23 | 10.3 (9-14) ^¶^ | | P | Case-control | EM, HFVM, nNO | - | - | + | + |
| BUSH et al. | 2006 | 19 | 9.5 ± 3.0^#^ | | P | Case-control | EM, HFVM, nNO | - | - | + | + |
| ZIHLIF et al. | 2005 | 20 | 10.8 (9-14) ^¶^ | | P | Case-control | EM, HFVM, nNO | - | - | + | - |
| CSOMA et al. | 2003 | 15 | 10.3 (7-14) ^*^ | | P | Case-control | EM, HFVM | - | - | + | - |
| **HORVATH et al.** | 2003 | 14 | 35±4.6^#^ | | P | Case-control | EM, HFVM | NA | - | + | + |
| NARANG et al. | 2002 | 31 | 11.0 (5.5-17.3) ^+^ | | P | Case-control | HFVM, EM | - | - | + | - |
| **KARADAG et al.** | **1999** | 21 | 10.8±3.2^#^ | | R | Case-control | EM, HFVM | NA | - | + | + |
| PHILLIPS et al. | 1998 | 12 | 11 (7-15) ^+^ | | P | Case-control | HFVM, EM | - | - | + | + |
| **USA** |  |  |  | |  |  |  |  |  |  |  |
| NOONE et al. | 1999 | 12 | 34 (14-71) ^+^ | | P | Case series | Clinical, EM | - (not by age) | - | + | + |
| REGNIS et al. | 2000 | 12 | 36 (13-70) ^+^ | | P | Case-control | Clinical, EM | - (not by age) | - | + | + |
| Age is reported in years as mean ±SD^#^, mean (range)^*^, median (IQR)^¶^, or median (range)^+^ depending on the available information; NR, not reported  R, retrospective; P, prospective; EM, electron microscopy; HFVM, high frequency video-microscopy; nNO, nasal nitric oxide; gen, genetic analysis; NA, not applicable  Studies appearing in bold are those included in the meta-analysis | | | | | | | | | | | |
| **Multi-centre/international studies** | | | |  | |  |  |  |  |  |  |
| **Denmark, Italy, UK** |  |  |  |  | |  |  |  |  |  |  |
| **MAGLIONE et al.** | 2014 | 158 | 8.7 (4.2–17.4) | R | | Case series | according to consensus | NA | + | + | + |
| **Netherlands, Italy** |  |  |  |  | |  |  |  |  |  |  |
| SANTAMARIA et al. | 2008 | 20 | 14.3 (4.6-27.5) ^+^ | P,R | | Case series | EM, LM | - | - | + | + |
| **Germany, Switzerland** |  |  |  |  | |  |  |  |  |  |  |
| **NYILAS et al.** | 2017 | 49 | 14.7±6.6^#^ | P | | Case-control | all tests | - | - | + | + |
| NYILAS et al. | 2016 | 20 | 13 ± 2.7^#^ | P | | Case-control | according to consensus | - | - | + | - |
| NYILAS et al. | 2015 | 47 | 14.7 ± 6.9^#^ | P | | Case-control | according to consensus | - | - | + | - |
| **Canada, USA** |  |  |  |  | |  |  |  |  |  |  |
| **DAVIS et al.** | 2015 | 118 | 8 (5-11)^¶^ | P | | Case series | EM or gen | NA | - | + | - |
| **NOONE et al.** | 2004 | 78 | adults: 36 (19-73) ^+^  children: 8 (1-17) ^+^ | P | | Case series | EM, HFVM, gen | + | - | + | - |
| **International PCD registry** |  |  |  |  | |  |  |  |  |  |  |
| WERNER et al. | 2016 | 201 | 20 (1-75) ^+^ | P | | Case series | all tests, including gen | - | + | + | + |

Age is reported in years as mean ±SD^#^, mean (range)*, median (IQR)^¶^, or median (range)^+^ depending on the available information; NR, not reported

R, retrospective; P, prospective; EM, electron microscopy; HFVM, high frequency video-microscopy; nNO, nasal nitric oxide; gen, genetic analysis; NA, not applicable

Studies appearing in bold are those included in the meta-analysis
