## Supplementary figures and images for "Spirometric indices in primary ciliary dyskinesia: systematic review and meta-analysis"

### Supplementary file 3

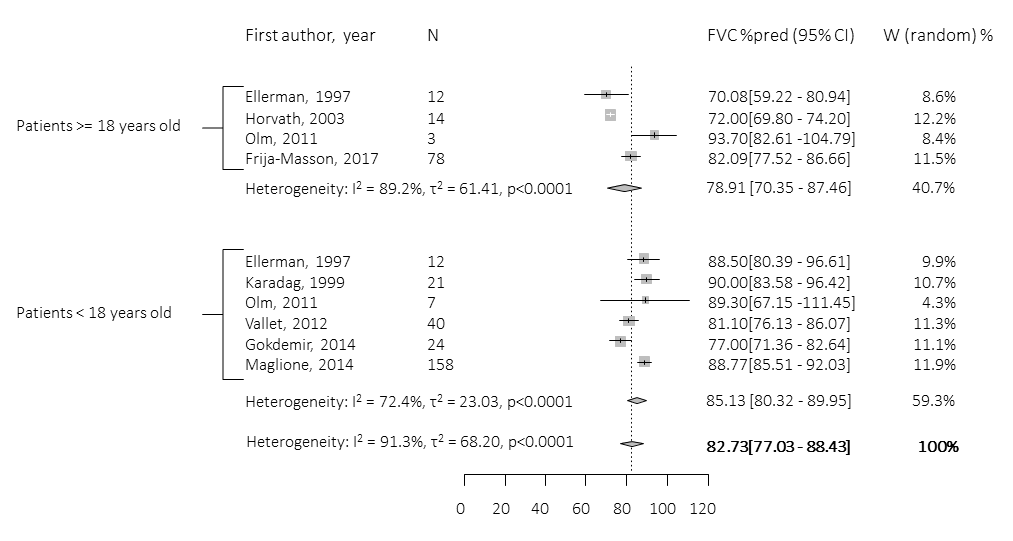
